# Inhibition of *agr* Quorum Sensing in Staphylococci by Novel Thiazole-Indole Hybrid Compounds

**DOI:** 10.1101/2025.10.15.682726

**Authors:** Weizhe Wang, Praveen Kumar Singh, Martin S. Bojer, Kirsi Savijoki, Jari Yli-Kauhaluoma, Hanne Ingmer, Jayendra Z. Patel, Stephanie Fulaz

**Author notes:** Corresponding authors on microbiology; on chemistry.

## Abstract

The rise of multidrug-resistant *Staphylococcus aureus* has prompted the search for innovative, non-bactericidal therapeutic strategies. Anti-virulence approaches targeting quorum sensing (QS) pathways offer a promising alternative by disarming pathogenicity, potentially without exerting selective pressure for resistance. In this study, we report the identification of two novel thiazole–indole hybrid compounds, namely 2-[(6-methoxy-1*H*-indol-2-yl)methyl]-5-cyano-1,3-thiazol-3-amine (designated **C3**) and 3-amino-5-cyano-2-(5-chloro-1*H*-indol-2-yl) thiazole (designated **C4**), as potent inhibitors of the *S. aureus* accessory gene regulator (*agr*) quorum sensing system. Building on two heterocyclic scaffolds commonly used in medicinal chemistry and associated with QS modulation, these synthetic small molecules act on the *S. aureus* AgrC QS receptor, attenuating virulence without affecting bacterial viability. Both compounds significantly reduced the production of secreted virulence factors without promoting biofilm formation, a known drawback of some other QS inhibitors. Furthermore, their activity extended to *Staphylococcus lugdunensis*, while sparing the commensal *Staphylococcus epidermidis*, indicating pathogen-selective QS interference. These findings establish novel thiazole–indole hybrids as a promising chemotype for anti-virulence intervention and provide valuable chemical tools for probing *agr*-mediated regulation in *Staphylococcus* species.

## Introduction

The escalating threat posed by antibiotic-resistant *Staphylococcus aureus*, including the methicillin-resistant (MRSA) strains, underscores the urgent need for alternative therapeutic strategies (Parmanik et al. 2022). Despite global efforts, the antibiotic development pipeline remains limited, and no effective vaccine is currently available (Septimus and Schweizer 2016; Clegg et al. 2021; Fait et al. 2024; Caldera et al. 2024; Tsai et al. 2022). In this context, anti-virulence therapies, which disarm rather than kill pathogens, have emerged as promising candidates for managing difficult-to-treat infections without imposing strong selective pressure for resistance development (Salam and Quave 2018).

A central regulatory system controlling virulence gene expression in *S. aureus* is quorum sensing (QS), mediated by the accessory gene regulator (*agr*) operon. Activation of *agr* is initiated by the *agrD*-encoded autoinducing peptides (AIPs) that bind to the sensor histidine kinase AgrC, leading to activation of the response regulator AgrA and subsequent upregulation of secreted virulence factors, including *α*-hemolysin encoded by *hla*, while cell-surface-attached virulence factors like Protein A encoded by *spa* are down-regulated (Yarwood and Schlievert 2003; Yamazaki et al. 2024). Interfering with this pathway through QS inhibitors (QSIs) has shown significant promise in preclinical studies (Daly et al. 2015; Sully et al. 2014; Todd et al. 2017; Baldry et al. 2018).

Several classes of natural and synthetic QSIs targeting different nodes in the staphylococcal *agr* pathway have been described, including AIPs from coagulase-negative staphylococci, cyclic peptides (e.g., solonamides), and cyclic lipopeptides such as fengycin, which act at the AgrC receptor (Nielsen et al. 2014; Piewngam et al. 2023; Piewngam et al. 2018; Brown et al. 2020; Paharik et al. 2017; Baldry et al. 2018). Other small molecules target AgrA (e.g., *ω*-hydroxyemodin, norlichexanthone, biaryl hydroxyketones, savarin, and 2-(4-methylphenyl)-1,3-thiazole-4-carboxylic acid) interfere with AIP biosynthesis via AgrB (e.g. ambuic acid) (Daly et al. 2015; Sully et al. 2014; Bezar et al. 2019; Khodaverdian et al. 2013; Todd et al. 2017; Nakayama et al. 2009; Baldry et al. 2016). However, a key challenge remains: *S. aureus* comprises four *agr* specificity groups (*agr* I-IV), which are often cross-inhibitory (Ji, Beavis, and Novick 1997), and many reported QSIs are effective only against select subgroups or lack molecular targets. Furthermore, some QSIs inadvertently promote biofilm formation or exhibit cytotoxicity (Choo, Rukayadi, and Hwang 2006; Bezar et al. 2019; Nielsen et al. 2014). Thus, the development of broad-spectrum QSIs with defined mechanisms of action remains a critical unmet need.

From a medicinal chemistry perspective, both thiazole and indole are privileged scaffolds with a wide spectrum of bioactivities. Thiazole derivatives are present in clinically approved antibiotics (e.g., penicillin), antitumor agents (e.g., tiazofurin), and antivirals (e.g., ritonavir), and have recently shown QS inhibitory effects (de Souza 2006; Althagafi, El-Metwaly, and Farghaly 2019; Ibrahim et al. 2020). For instance, 5-acetyl-4-methyl-2-(3-pyridyl) thiazole suppresses hemolysin and biofilm production in MRSA and *Chromobacterium violaceum* (Ibrahim et al. 2020). Similarly, the indole core is a fundamental motif in numerous biologically active compounds, including serotonin, melatonin, and various antimicrobial and anticancer agents (Abdellatif, Lamie, and Omar 2016; Kaur, Singh, and Narasimhan 2019). Beyond its pharmaceutical relevance, indole functions as a microbial signalling molecule influencing QS behavior across species (Lee et al. 2013; Tao et al. 2023).

Molecular hybridization, merging two bioactive fragments into a single structural framework, is a well-established strategy to enhance potency, broaden activity profiles, and minimize resistance development (Ivasiv et al. 2019; Simakov et al. 2021). Guided by this rationale, we hypothesized that integrating thiazole and indole pharmacophores, whose derivatives have been reported as QS modulators, could enhance *S. aureus* QS inhibition. To explore this hypothesis, we synthesized and screened a focused library comprising both thiazole derivatives and thiazole–indole hybrids for QS inhibitory activity.

Preliminary screening identified several compounds with QS-related inhibitory activity in *S. aureus*. Two lead molecules, **C3** and **C4**, demonstrated broad-spectrum *agr* inhibition of all four *S. aureus agr* groups, likely through competitive binding to the AgrC receptor. These compounds effectively suppressed RNAIII-driven virulence factor production and reduced hemolytic activity without promoting biofilm formation. Furthermore, their activity extended to *Staphylococcus lugdunensis*, a pathogenic coagulase-negative species in which the *agr* quorum-sensing system contributes to the regulation of virulence gene expression (Chin et al. 2022), while sparring the skin commensal *S. epidermidis*. Taken together, our findings position thiazole-indole hybrids as promising scaffolds for anti-virulence therapy targeting quorum sensing in staphylococci.

## Materials and Methods

### Bacterial strains and growth conditions

*S. aureus, S. epidermidis, S. lugdunensis*, and *S. schleiferi* strains were grown in tryptone soya broth (TSB) media (Oxoid), at 37 °C with shaking at 200 rpm. When required, relevant antibiotics (Sigma-Aldrich) were added at the following final concentrations: 10 μg/mL chloramphenicol or 5 μg/mL erythromycin for *S. aureus, S. epidermidis*, and *S. lugdunensis*, respectively.

### General synthetic scheme of the compounds (C1–C5)

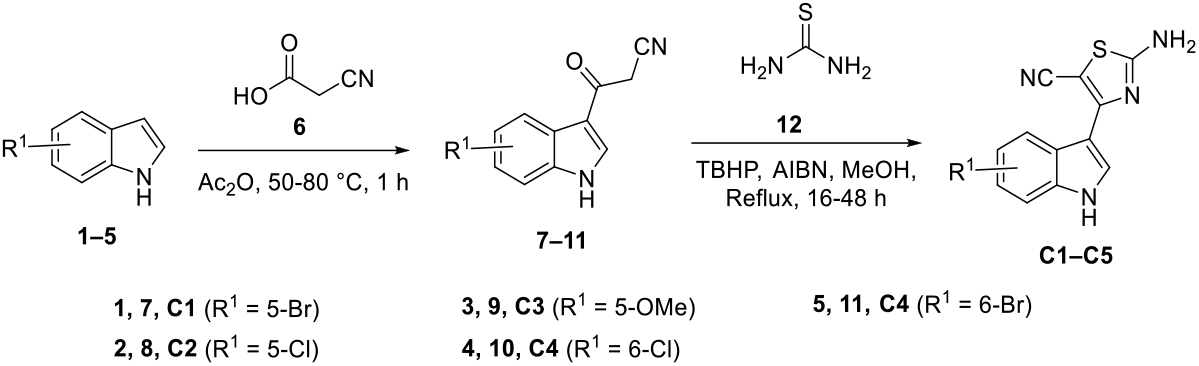

The thiazole compound library was synthesized (see general synthetic scheme and supporting information) at the University of Helsinki as a part of our ongoing research. The novel thiazole–indole hybrid compounds were designed based on the meridianin D in which 6-membered heterocyclic scaffold pyrimidine was replaced by 5-membered heterocyclic scaffold thiazole to attain novelty. A small set of thiazole–indole hybrid compounds, such as **C1**–**C5** were synthesized conveniently using commercially available starting materials (see general synthetic scheme). First, commercially available indoles **1**–**5** were subjected to acylation using cyanoacetic acid **6** in presence of acetic anhydride to give intermediates **7**–**11**. The obtained acylated indole intermediates **7**–**11** were subjected to cyclization using thiourea **12** under reflux conditions to yield the targeted thiazole–indole hybrid compounds.

### Reporter assays

A qualitative agar plate assay was performed to monitor expression of *spa* gene using the *spa-lacZ* reporter fusion strain PC203 as described previously (Bojer, Baldry, and Ingmer 2017; Nielsen et al. 2010).

For liquid assays, *β*-lactamase reporter tests were performed as previously described with modifications (Gless et al. 2019), using RN10829 AgrC WT (P2-*agrA*; P3-*blaZ*)/p*agrC*-*I*-*WT* and RN10829 AgrC const. (P2-*agrA*; P3-*blaZ*)/pEG11(*agrC*-*I*-*R238H*), and exposed to supernatants from *S. aureus* 8325-4 in the presence of **C3** (200 µM, 100 µM), **C4** (50 µM, 25 µM), or DMSO. *β*-lactamase activity was assessed by colorimetric measurement of nitrocefin hydrolysis, and the rate of hydrolysis (reaction slope) was used as a quantitative readout of *agr* system activation.

Fluorescent reporter assays were performed as previously described (Gless et al. 2025; Wang et al. 2025) using *S. aureus agr I-IV* (P3-*yfp*), *S. epidermidis agrI-III* (P3-*gfp*), and *S. lugdunensis agrI* (P3-*gfp*) reporter strains. Cells were exposed to **C3** or **C4** at the following concentration ranges: –for *S. aureus*: **C3** (200–6.25 µM), **C4** (50–3.125 µM); – for *S. epidermidis* and *S. lugdunensis*: **C3** (200–50 µM), **C4** (50–12.5 µM). DMSO was included as a vehicle control. Promoter activity was monitored over 24 h at 37 °C under continuous orbital shaking using YFP or GFP fluorescence, measured with a Synergy H1 microplate reader (BioTek).

### Dose-response analysis and quality assessment

The IC_50_ values were calculated using nonlinear regression with a four-parameter logistic model (4PL) in OriginPro 2020 (OriginLab, Northampton, MA, USA). Results are expressed as mean ± SEM unless otherwise indicated.

For assay quality evaluation, Z′-factors were calculated in Microsoft Excel based on the mean and standard deviation of the positive and negative controls, using the formula:

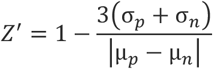

where *μ*_*p*_ and *μ*_*n*_ are the means, and *σ*_*p*_ and *σ*_*n*_ are the standard deviations of the positive and negative controls, respectively. Z′ value ≥ 0.5 indicates an excellent assay; values between 0 and 0.5 are considered marginal, and values < 0 are unacceptable (Zhang, Chung, and Oldenburg 1999).

### Hemolytic activity

Hemolytic activity was determined by measuring blood cell lysis as previously described (Diaz et al. 2024; Wang et al. 2025), using culture supernatants from *S. aureus* 8325-4 grown with C3 (200–50 µM) or C4 (50–12.5 µM) at an initial OD_600_ of 0.02. Cultures treated with DMSO served as the control. In parallel, the intrinsic hemolytic activity of **C3** and **C4** was assessed by incubating the compounds directly with red blood cells in the absence of bacteria.

### Biofilm formation assay

Quantitative biofilm formation assays were performed as previously described (Bezar et al. 2019), using *S. aureus* 8325-4 cultures treated with **C3, C4**, or DMSO. After 24 hours static incubation in 96-well plates, biofilm biomass was quantified using crystal violet staining, followed by measurement of absorbance at 595 nm with a Synergy H1 microplate reader (BioTek).

## Results

### Thiazole–Indole Hybrids Selectively Inhibit the *agr* QS System in *S. aureus*

To search for compounds inhibiting the *S. aureus agr* quorum sensing system, we screened a small compound library containing thirteen thiazole and five thiazole-indole hybrid compounds (Figure 1A; Table S1). These compounds were evaluated using an *S. aureus agrI* P3-*yfp* reporter strain, in which the promoter of RNAIII (P3) – a key effector of the *agr* operon - is fused to *yfp* encoding the yellow fluorescence protein (YFP) gene. Among the 18 compounds tested, four showed no measurable effect, and the remaining 14 demonstrated varying degrees of inhibition without affecting bacterial growth (Figure 1A, S1, and S2). Of these inhibitors, **C3** and **C4** stood out by retaining activity at lower concentrations and showing concentration-dependent suppression of YFP fluorescence. To further validate their specificity toward *S. aureus agr*, we assessed *spa* expression using the PC203 *spa-lacZ* reporter strain. The *spa* gene, which encodes protein A, is negatively regulated by RNAIII; thus, inhibition of *agr* derepresses *spa* transcription. Consistent with RNAIII inhibition, **C3** and **C4** elevated *β*-galactosidase activity expressed from the *spa-lacZ* fusion (Figure 1B). Together, these results demonstrate that **C3** and **C4** act selectively on the QS circuitry rather than affecting growth.

**Figure 1.**
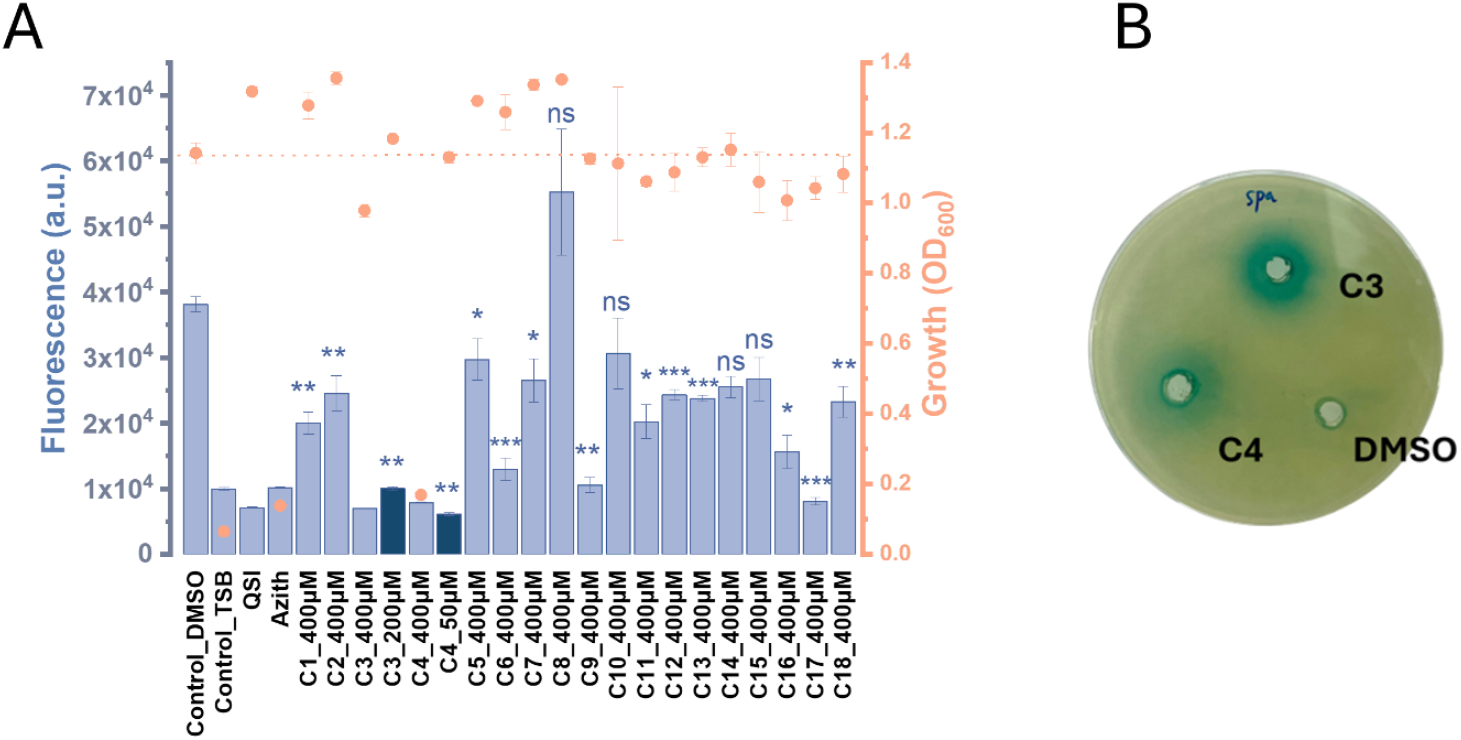
The inhibitory effect of test compounds on *S. aureus agr* activity. (**A**) *S. aureus agrI* P3-*yfp* reporter strain was cultured in the presence of the indicated concentrations of compounds C1–C18. After 24 h, optical density at 600 nm (OD_600_, orange dots) and fluorescence intensity (YFP signal, blue bars) were measured. Compounds were tested at 400 µM (blue) and at lower concentrations - 200, 100, 50, and 25 µM - shown in dark blue. A supernatant from *S. schleiferi* was used as a QSI positive control, and azithromycin (Azith) served as a bacterial growth inhibition control. Data are shown from a representative biological replicate, each test measured in duplicate. (**B**) Effect of compounds **C3** and **C4** on *spa* expression was monitored by adding 15 µL of **C3** and **C4** (20 mM) to wells in TSA plates containing the *S. aureus* 8325-4 derived *lacZ* reporter strain PC203 (*spa-lacZ*), erythromycin and X-gal.

### Compounds Attenuate *α*-Hemolysin-Mediated Hemolysis

To assess the functional impact of *agr* inhibition on toxin production, we measured the hemolytic activity of culture supernatants from *S. aureus* strains treated with compounds **C3** and **C4**. Hemolysis in *S. aureus* is primarily mediated by *α*-hemolysin (Hla), a major virulence factor positively regulated by the *agr* QS system. As expected, treatment with either compound led to a marked, dose-dependent decrease in hemolytic activity, achieving ∼90% reduction at the highest tested concentrations (Figure 2).

**Figure 2.**
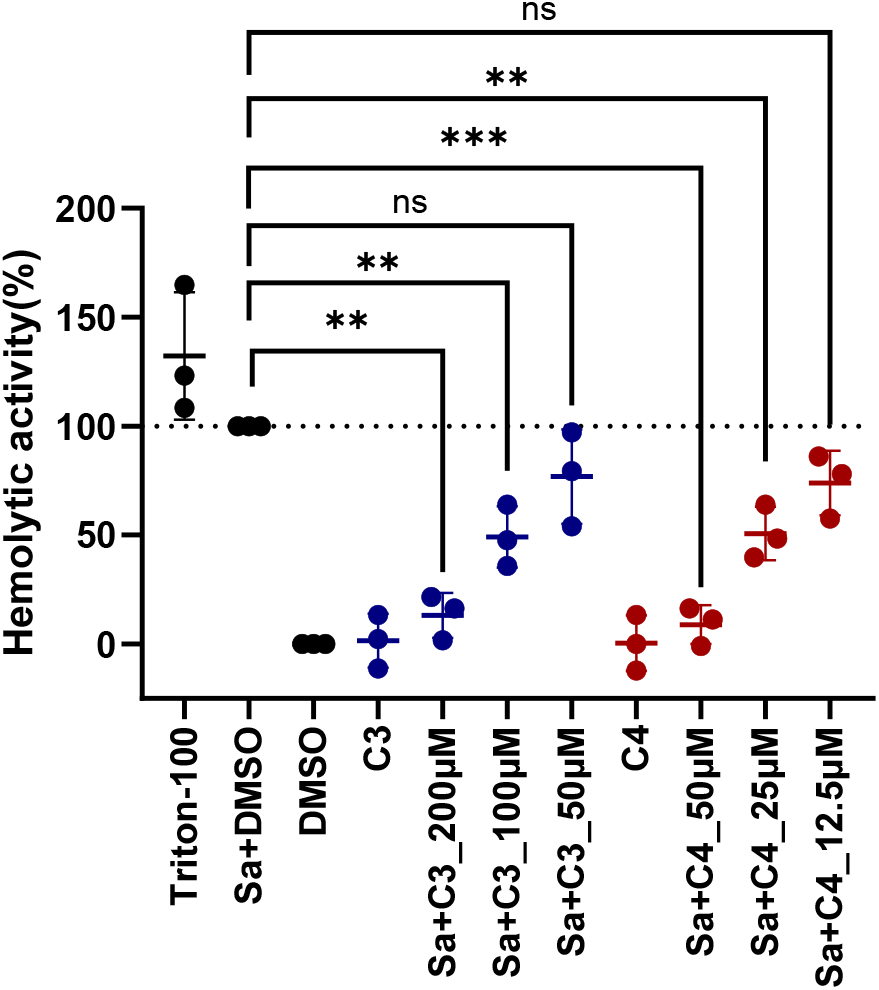
C3 and C4 modulation of *S. aureus* hemolytic activity. Hemolysis was measured using the highly hemolytic strain, *S. aureus* 8325-4 (Sa) with or without the testing compounds. After overnight incubation supernatants were collected, incubated with bovine blood cells for one hour, centrifuged, and absorbance was measured at OD_450_. Hemolytic activity was normalized to the full hemolysis control (*S. aureus* cultured with DMSO) and the background control (TSB medium with DMSO). Data are represented as mean ± SD of at least 3 technical replicates from a representative biological replicate. Statistical significance was determined by Student’s t-test compared to the full hemolysis control. Statistical marker ** (p < 0.01) and *** (p < 0.001) highlights significant differences in activity levels compared to the control. ns, no significant difference.

Importantly, neither **C3** nor **C4** displayed intrinsic hemolytic activity against human erythrocytes, showing that the observed reduction in hemolysis is not due to compound-mediated disruption of host membranes. This supports a non-cytolytic, target-specific mode of action consistent with inhibition of the *agr* system by **C3** and **C4** effectively diminishing virulence factor production.

### The AgrC Sensor Kinase as the Molecular Target

To investigate the mode of action, we examined the activity of **C3** and **C4** in a reporter strain that expresses a constitutively active AgrC variant (R238H), which bypasses AIP dependency (Geisinger, Muir, and Novick 2009). This system allows us to determine whether a compound acts upstream or downstream of AgrC, as only downstream-acting compounds are expected to inhibit RNAIII expression in the constitutively active variant. Indeed, both **C3** and **C4** reduced RNAIII expression in the reporter strain expressing the WT AgrC (RN10829 AgrC WT) but had no significant effect on the AgrC constitutively active strain (RN10829 AgrC const., Figure 3, Table S2), suggesting that both compounds inhibit *agr* by competitively binding or otherwise interfering with AgrC activation.

**Figure 3.**
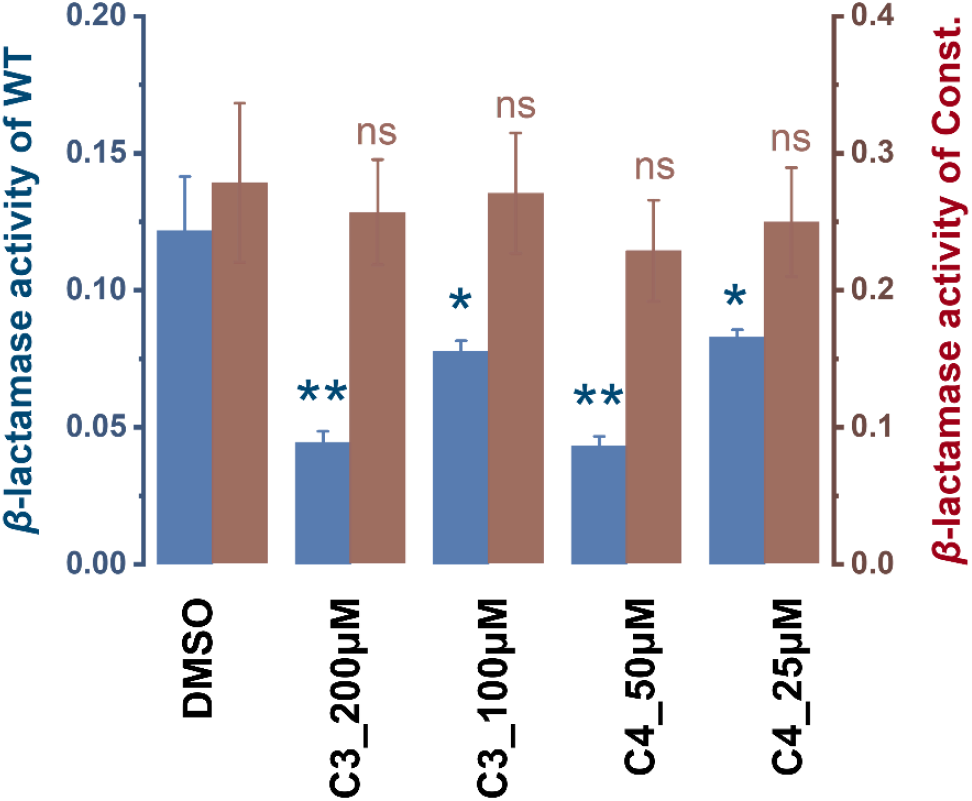
C3 and C4 do not influence RNAIII expression in the presence of a constitutively active AgrC. Cultures of RN10829 AgrC WT (blue) and RN10829 AgrC constitutively active (brown) grew to OD_600_ 0.35 where 1/10 a volume of *S. aureus* 8325-4 supernatant and testing compounds or DMSO was added. Samples were obtained 30 min after addition and were analyzed for *β*-lactamase activity (RNAIII expression). Data are shown from a representative biological replicate, each test measured in three technical replicates.. *β*-lactamase activity was quantified by measuring nitrocefin hydrolysis rates (reaction slopes). Statistical significance was determined by comparison to the control group using student’s *t*-test. ns (p > 0.05) represents no significant difference between test conditions; * (p < 0.05) and ** (p < 0.01) represent a significant difference between test conditions.

### Broad-Spectrum Activity across *S. aureus agr* Specificity Groups

The *agr* locus in *S. aureus* exists in four polymorphic variants (*agr I–IV*), each encoding a distinct AIP with group-specific receptor interactions. Using P3-*yfp* reporter strains representing each *agr* group, we examined the effects of **C3** and **C4** over a range of concentrations (Figures S1–S3). We observed that both compounds inhibited RNAIII expression in a concentration-dependent manner across all four *agr* groups. Half-maximal inhibitory concentrations (IC_50_) were calculated using logistic regression, considering only concentrations that did not affect growth (Figure S3). **C4** consistently demonstrated greater potency, with IC_50_ values ranging from 7 ± 1 μM to 26 ± 3 μM across groups *I–IV* (Table 2). **C3** was less potent overall, with IC_50_ values spanning 22 ± 2 μM to 80 ± 5 μM. The lowest IC_50_ values were observed against *agrII*, while *agrIII* exhibited the least sensitivity to the compound, highlighting a potential influence of AgrC sequence divergence on compound affinity.

**Table 1.**
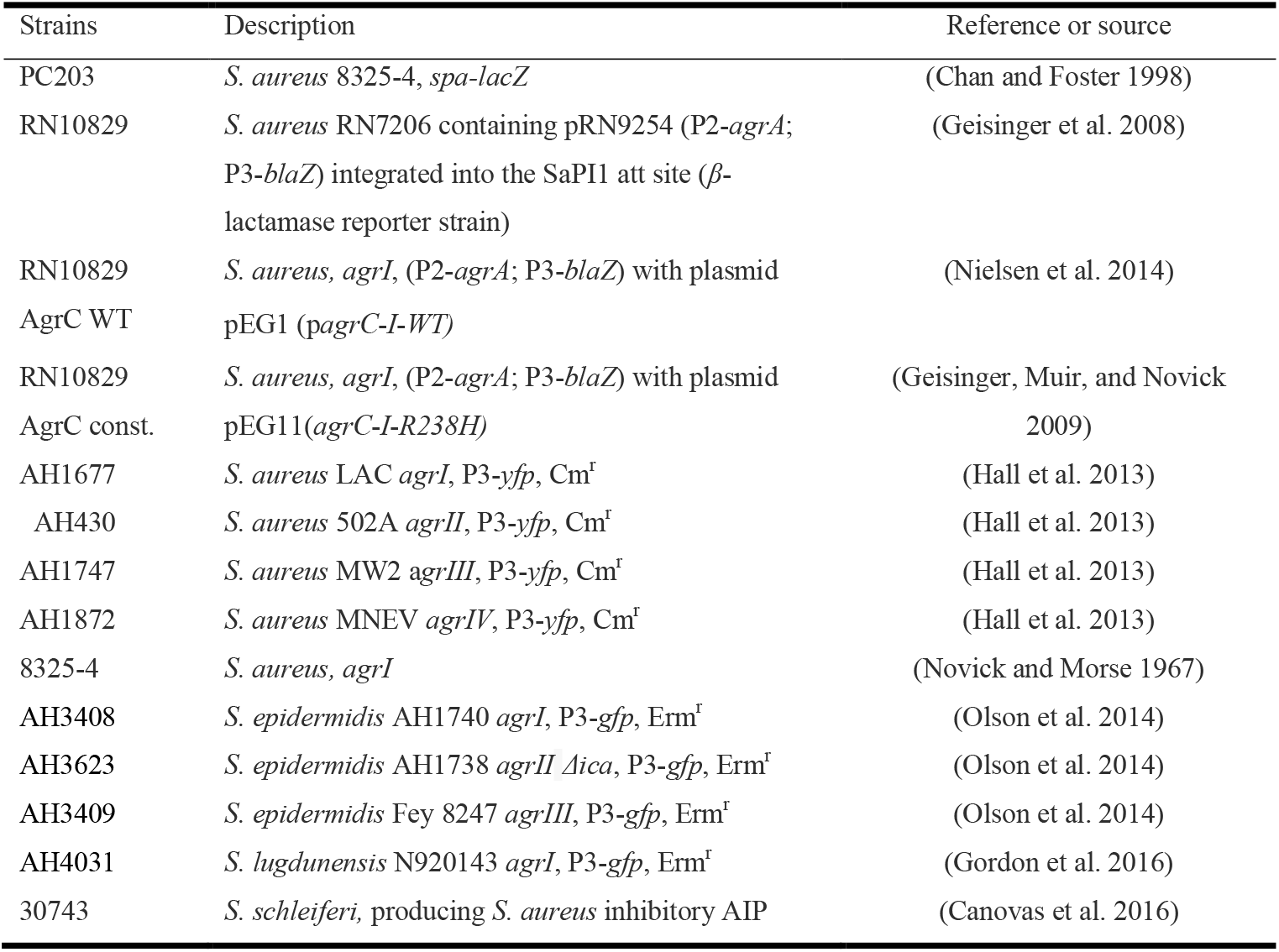
Bacterial strains used in this study.

**Table 2.** The half-maximal inhibitory concentration (IC_50_) values of compounds **C3** and **C4** against *S. aureus agr I-IV*.

| IC<sub>50</sub> ( μM ) | *agrI* | *agrII* | *agrIII* | *agrIV* |
| --- | --- | --- | --- | --- |
| **C3** | 41±3 | 22±2 | 80±5 | 43±6 |
| **C4** | 8±1 | 7±1 | 26±3 | 18±4 |

These findings demonstrate that both **C3** and **C4** achieve broad-spectrum inhibition of *S. aureus agr* subtypes, a critical feature for therapeutic translation given the prevalence of *agr* polymorphisms in clinical isolates.

### Differential Modulation of *agr* in Coagulase-negative Staphylococci

To evaluate whether **C3** and **C4** inhibit *agr* in other staphylococcal species, we tested **C3** and **C4** against P3-*gfp* reporter strains of *S. epidermidis* (*agr I–III*) and *S. lugdunensis* (*agr I*). These species also utilize AIP-AgrC signalling, but their AgrC sequences diverge from those in *S. aureus*, potentially influencing compound responsiveness (Thoendel et al. 2011).

In *S. epidermidis*, **C3** inhibited at high concentrations *agr* in a strain carrying *agrI*, while neither compound affected *agr* in a strain carrying *agrIII*. Unexpectedly, both compounds enhanced RNAIII expression in the *S. epidermidis agrII* strain. Also this inducing effect was seen even in the presence of an AIPI from *S. simulans* that previously was shown to be inhibitory to *S. epidermidis agrII* (Figure 4, S6–S7) (Gless et al. 2025). This suggests agonist-like behaviour and potential non-competitive binding to AgrC. In contrast, both compounds significantly suppressed *agr* activity in *S. lugdunensis*, including at low micromolar concentrations (Figure 4, S4–S5).

**Figure 4.**
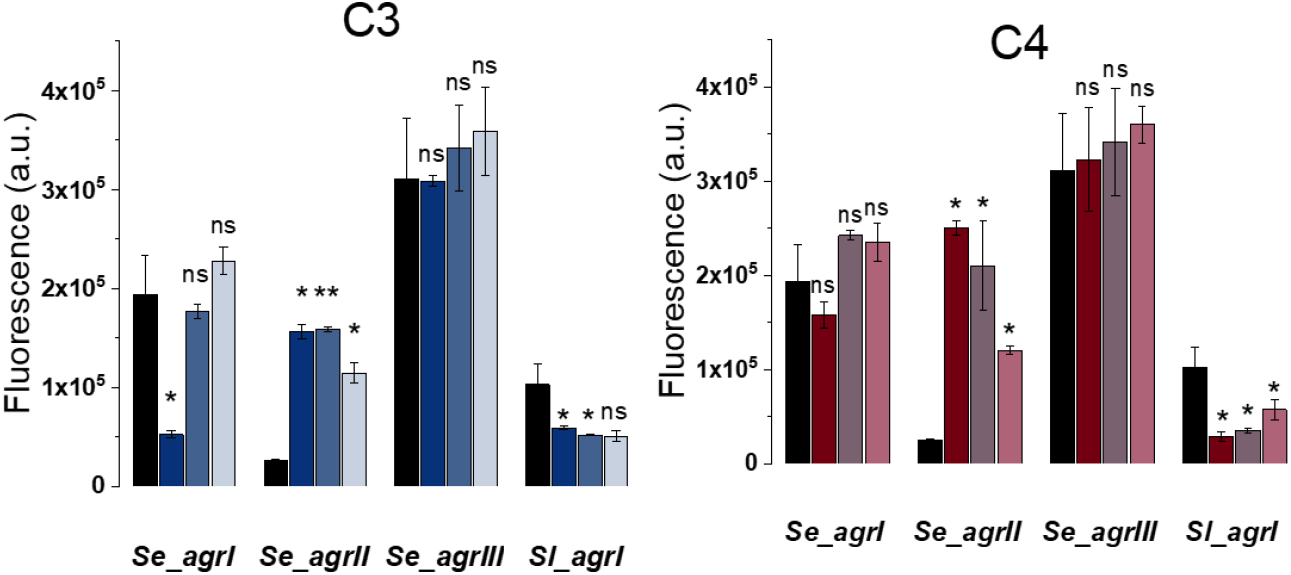
Influence of C3 and C4 on *S. epidermidis agrI-III*, and *S. lugdunensis agrI* QS activity. P3 promoter activity was monitored over 24 hours using *S. epidermidis* (Se) P3-*gfp* reporter strains carrying *agrI, agrII* and *agrIII* alleles, respectively, and a *S. lugdunensis* (Sl) P3-*gfp* reporter strains in the presence or absence of test compounds. Statistical significance was determined using a student’s *t*-test by comparing test conditions: ns (p > 0.05) indicates no significant difference, * (p ≤ 0.05) and ** (p ≤ 0.01) indicate significant differences. Black: untreated GFP reporter strains; Blue: GFP reporter strains treated with **C3** (deep blue: 200 µM; medium blue: 100 µM; light blue: 50 µM); Red: GFP reporter strains treated with **C4** (deep red: 50 µM; medium red: 25 µM; light red: 12.5 µM).

These results underscore the context-dependence of *agr* modulation and highlight that interactions between thiazole–indole hybrids and AgrC are likely influenced by species-specific receptor architecture. Selective inhibition of pathogenic species (*S. aureus, S. lugdunensis*) while sparing commensals (*S. epidermidis*) represents an ideal profile for anti-virulence agents.

### C3 and C4 do not Impact Biofilm Formation

The *agr* system plays a complex role in *S. aureus* biofilm formation, influencing development either negatively or positively depending on the stage (Yamazaki et al. 2024; Schilcher and Horswill 2020). Inactivation of *agr* in *S. aureus* has been linked to enhanced biofilm formation under various conditions (Bezar et al. 2019; Dotto et al. 2021; Vuong et al. 2000). Given this, we evaluated the effects of **C3** and **C4** on biofilm development in *S. aureus* 8325-4 using the crystal violet assay. Neither compound significantly altered biomass accumulation compared to DMSO controls (Figure 5). This indicates that, under the tested conditions, *agr* inhibition by **C3** and **C4** does not promote biofilm formation.

**Figure 5.**
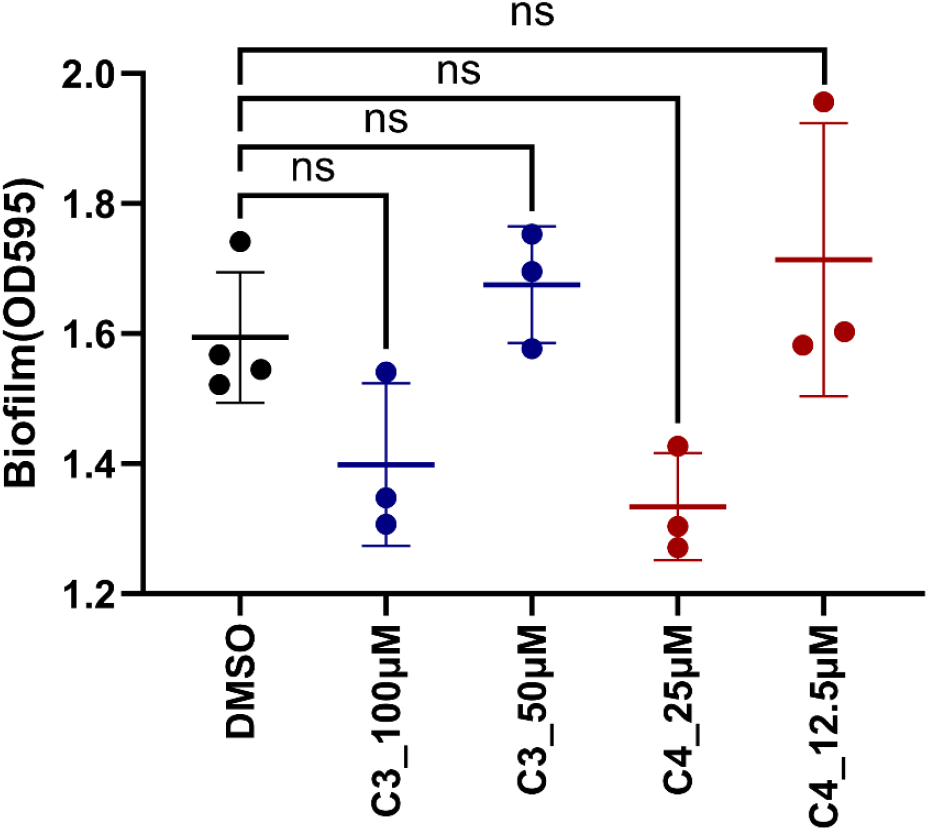
C3 and C4 do not affect biofilm formation. *S. aureus* 8325-4 culture was cultivated in TSB media in the absence or presence of the testing compounds. The quantification of the biomass was performed after 24 h using the crystal violet assay, and absorbance was measured at OD595. Data represented as mean ± SD of at least 3 technical replicates from a representative biological replicate. Statistical significance was determined using Student’s t-test; ns (p > 0.05) represents no significant difference between test conditions.

### Impact of substituents on thiazole-indole scaffold

Among **C1**–**C5** compounds, subtle changes to the indole substituents dramatically affected potency. **C3** (5-methoxy) and **C4** (6-chloro) emerged as the most active compounds, whereas brominated analogs (**C1** having 5-bromo while **C5** having 6-bromo) were considerably less effective. Interestingly, the shifting of chloro from 6^th^ (i.e., **C4**) to 5^th^ position (i.e., **C2**) also resulted in less effective compounds. This suggests that both electronic effects and steric factors at the 5^th^ and 6^th^ positions of the indole ring play a critical role in modulating interaction with AgrC. Such kind of activity modulation was seen depending on positions of different substituents on indole QS inhibitors (Lee et al. 2013).

The hybridization of thiazole and indole scaffolds into a single pharmacophore likely contributes to dual-site interactions at the receptor interface. This design principle is commonly employed in rational drug design to enhance target affinity, stability, and selectivity.

## Discussion

Throughout the antibiotic era, antibiotic resistance has arisen and spread quickly, often within a few years after the introduction of an antibiotic class to clinical use (Dickey, Cheung, and Otto 2017). Over the past 15 years, there has been a consistent and substantial increase in the use of antibiotics, a trend further exacerbated by the prolonged SARS-CoV-2 pandemic (Lázár et al. 2022). Recognizing the limitations of traditional antibiotics, the FDA has endorsed the development of anti-virulence therapies under specific regulatory frameworks. A landmark example is the approval of bezlotoxumab (Zinplava), a monoclonal antibody targeting *Clostridioides difficile* toxin B (Lowy, Molrine, and Ambrosino 2010).

Among bacterial pathogens, *S. aureus* warrants particular attention due to its formidable resistance against multiple antibiotic classes and its ability to cause a wide range of infections, from skin abscesses to life-threatening sepsis (Tong et al. 2015). Moreover, the epidemiological success of community-associated MRSA strains is due in part to increased virulence, which is driven by the presence and enhanced production of multiple immune evasion molecules and toxins (Otto 2010). Many virulence factors of *S. aureus* offer opportunities for anti-virulence intervention. For instance, methylophiopogonanone A (MoA) has been identified as an inhibitor of MgrA, a pivotal global regulator of virulence (Guo et al. 2024). Additionally, 3,6-disubstituted triazolothiadiazole compounds have demonstrated inhibitory activity against sortase A (SrtA), a membrane-anchored transpeptidase responsible for anchoring surface adherence and immune evasion proteins to the bacterial cell wall (Zhang et al. 2014). The *agr* QS system, as a master regulator of many staphylococcal virulence factors, continues to be an especially attractive anti-virulence target.

Here, we report two novel thiazole–indole hybrid compounds as potent inhibitors of the *S. aureus agr* QS system, namely 2-[(6-methoxy-1*H*-indol-2-yl) methyl]-5-cyano-1,3-thiazol-3-amine (**C3**) and 3-amino-5-cyano-2-(5-chloro-1*H*-indol-2-yl) thiazole (**C4**). These hybrids act via competitive inhibition of the membrane-bound sensor kinase AgrC, a mechanism supported by their lack of activity in strains expressing a constitutively active AgrC variant. Both compounds demonstrated broad-spectrum inhibition across all four *agr* specificity groups, with **C4** exhibiting greater potency (IC_50_ 7–26 µM) than **C3** (IC_50_ 22–80 µM), while sparing bacterial viability and preserving growth.

From a medicinal chemistry perspective, the thiazole–indole scaffold offers a privileged structural motif with multiple pharmacophoric vectors. The thiazole ring contributes to π-stacking, metal coordination, and hydrogen bonding (Chen et al. 2025; Rossin et al. 2011), while the indole moiety serves as a planar, hydrogen-bond-rich anchor commonly found in kinase inhibitors (Zhang et al. 2023), G protein-coupled receptor (GPCR) ligands (Sabnis 2025), and bacterial signal disruptors (Lee et al. 2013). Our results revealed that subtle electronic and steric modifications at the indole *C*-5 and *C*-6 positions significantly influenced inhibitory activity. The methoxy substituent at *C*-6 in **C3** supported moderate activity, whereas the *C*-5 chlorine in **C4** enhanced potency, likely through improved electronic complementarity or hydrophobic interactions with the AgrC binding pocket. Brominated analogues (**C1, C5**), however, exhibited diminished activity, suggesting that excess steric bulk at these positions is detrimental to binding.

These observations align with prior reports on indole-based QSIs, such as 7-benzyloxyindole (7BOI), which influence virulence gene expression without affecting bacterial viability (Lee et al. 2013). Additionally, the thiazole–indole motif has been shown to possess antimicrobial activity through alternative mechanisms, suggesting potential for dual-function scaffolds (Simakov et al. 2021). Together, these findings position this hybrid chemotype as a promising platform for further optimization.

A key challenge in QS inhibition lies in achieving species and strain specificity. Our data demonstrate that **C3** and **C4** suppress *agr* activity in *S. aureus* and the closely related pathogen *S. lugdunensis*, but not in the skin commensal *S. epidermidis*, where the compounds were either inactive or—interestingly, in the case of *agrII*—weakly agonistic. This may indicate direct activation of AgrC, although further studies are needed. This selective inhibition pattern mirrors that of savirin (Sully et al. 2014), which targets *S. aureus* while sparing commensals. Such precision is desirable, as it minimizes off-target effects on beneficial microbiota and reduces the likelihood of dysbiosis.

Regulation of biofilm formation presents another layer of complexity in *S. aureus* pathogenesis. *agr*-mediated QS modulates biofilm dispersal, yet its inhibition has been paradoxically associated with both increased and decreased biofilm formation depending on the environmental context (Schilcher and Horswill 2020). Importantly, neither **C3** nor **C4** significantly altered biofilm biomass under static growth conditions, suggesting that inhibition of *agr* by these compounds does not promote biofilm accumulation under the tested conditions.

Taken together, our findings highlight thiazole–indole hybrids as chemically tractable anti-virulence agents that achieve broad *agr* inhibition, attenuate virulence outputs such as hemolytic activity, and exhibit species-level selectivity without impacting bacterial growth or biofilm development. By acting through a non-bactericidal mechanism and targeting a non-essential but central regulatory system, these compounds exemplify the principles of “pathoblocker” design, selective disarmament of pathogens without ecological disruption.

## Acknowledgements

We thank Prof. Christian A. Olsen and Postdoc Benjamin S. Bejder (Department of Drug Design and Pharmacology, University of Copenhagen) for their valuable guidance, access to infrastructure, and for providing the synthesized autoinducing peptides (AIPs). Funding was provided by LEO Foundation: LF-OC-20-000517 and Novo Nordisk Foundation: NNF22OC0077593 as well as by the Chinese Scholarship Council to WW. PKS would like to thank the Finnish Cultural foundation (grant numbers 00211014, 00221167, and 00231288) while JZP would like to thank the Academy of Finland (grant number 322724), which supported this work.

